# Structural studies reveal that endosomal cations promote formation of infectious CVA9 A particles, facilitating RNA and VP4 release

**DOI:** 10.1101/2022.09.02.506448

**Authors:** Aušra Domanska, Zlatka Plavec, Visa Ruokolainen, Benita Löflund, Varpu Marjomäki, Sarah J Butcher

**Affiliations:** Faculty of Biological and Environmental Sciences, Molecular and Integrative Bioscience Research Programme, and Helsinki Institute of Life Sciences-Institute of Biotechnology, University of Helsinki, Helsinki, Finland; Department of Biological and Environmental Sciences, Nanoscience Center, University of Jyväskylä, Finland

## Abstract

Coxsackievirus A9, an enterovirus, is a common cause of paediatric aseptic meningitis and neonatal sepsis. During cell entry, enterovirus capsids undergo conformational changes leading to expansion, formation of large pores, externalization of VP1 N-termini and loss of the lipid factor from VP1. Factors such as receptor binding, heat, and acidic pH can trigger capsid expansion in some enteroviruses. Here we show that fatty-acid free bovine serum albumin or neutral endosomal ionic conditions can independently prime CVA9 for expansion and genome release. Our results show that CVA9 treatment with albumin or endosomal ions generates a heterogeneous population of virions, which could be physically separated by asymmetric flow field flow fractionation and computationally by cryo-EM and image processing. We report cryo-EM structures of CVA9 A-particles obtained by albumin or endosomal ion treatment and a control non-expanded virion to 3.5, 3.3 and 2.9 Å resolutions, respectively. Where albumin promotes stabile expanded virions, the endosomal ionic concentrations induce unstable CVA9 virions which easily disintegrate losing their genome. Loss of most of the VP4 molecules and exposure of negatively-charged amino acid residues in the capsid’s interior after expansion, create a repulsive viral RNA-capsid interface, aiding genome release.

**Importance:** Coxsackievirus A9 (CVA9) is a common cause of meningitis and neonatal sepsis. The triggers and mode of action of RNA release into the cell unusually do not require receptor interaction. Rather, a slow process in the endosome, independent of low pH is required. Here, we show by biophysical separation, cryogenic electron microscopy and image reconstruction that albumin and buffers mimicking the endosomal ion composition can separately and together expand and prime CVA9 for uncoating. Furthermore, we show in these expanded particles that VP4 is present at only ~10% of the occupancy found in the virion, VP1 is externalised and the genome is repelled by the negatively-charged, repulsive inner surface of the capsid that occurs due to the expansion. Thus, we can now link observations from cell biology of infection with the physical processes that occur in the capsid to promote genome uncoating.

## Introduction

Coxsackievirus A9 (CVA9) belongs to the *Enterovirus* genus, *Enterovirus B* species (EV-B) in the family *Picornaviridae*. CVA9 can frequently cause paediatric aseptic meningitis and neonatal sepsis (1–4). Like other picornaviruses, CVA9 has a small, ~30 nm in diameter, non-enveloped viral capsid. Its (+)ssRNA genome is ~7 500 bases long and encodes for 4 structural (VP1-4) and 7 non-structural proteins (2A-C, 3A-D) (5). Expanded A-particles are described for many enteroviruses, occurring at early steps during virus entry (6–10). It is thought that VP4 must be present in the A-particle to be infectious, but VP4 is very difficult to detect (11). Many picornaviruses, including members of the EV-B, accommodate a lipid factor in the VP1 hydrophobic pocket. Palmitate is a commonly found lipid factor/fatty acid in enteroviruses (12). Expulsion of the lipid factor is associated with virion expansion. Once expansion has occurred, and the capsid has been endocytosed, genome release can occur (13). It has been shown that receptor binding can trigger a loss of the lipid factor and virion expansion in some of the EV-B members (14), but not in CVA9 and echovirus 1 (E1) (15–17). In addition, CVA9 and E1 do not depend on low pH for genome release, unlike members of *Enterovirus A* species indicating that these viruses use additional cues for uncoating (15, 18). Despite extensive research, the physiological triggers leading to enterovirus uncoating after internalization, other than receptor binding and low pH in the endosomes, are poorly understood (19). Some reports show serum albumin as an uncoating cue for E1, E12 and coxsackievirus B3 (6, 20, 21).

It is well established that acidification is key to endosome maturation. In addition to proton influx during endosome maturation, the concentrations of other ions in the endosomal lumen such as sodium, potassium and calcium also change over time due to ion channels and pumps present in the endosomal membranes (22, 23). The absolute values of ion concentrations within the endosome structures are not known, and they can notably vary even within specific microenvironments of the endosome (22, 23). However, the general trend is a decrease in sodium and calcium concentrations and an increase in potassium concentration during the endosome maturation (24–26). Thus, changes in ion concentrations other than protons may well serve as a trigger for virus uncoating.

We showed previously that, at 37 °C, albumin in combination with a neutral buffer mimicking the endosomal environment triggers the expansion of E1 without receptor engagement creating an infectious A-particle (6). However, we did not show the individual effect of either albumin or ion changes to the overall conformational changes identified. Here we show by real-time spectroscopy, gradient ultracentrifugation, cryo-EM and image reconstruction that endosomal ionic concentrations alone are sufficient to induce CVA9 expansion at a physiologically-relevant temperature. Moreover, this treatment leads to unstable CVA9 virions which easily disintegrate losing their genome. Albumin alone or in combination with endosomal ionic concentrations induces CVA9 expansion. We also show that particle expansion is a dynamic process yielding a mixed population of virions, as assessed by gradient ultracentrifugation and cryo-EM analysis. Using asymmetric flow field flow fractionation (AF4), we were able to physically separate expanded and intact particles and to confirm by mass spectrometry (MS) that expanded particles largely lack the small internal capsid protein, VP4. Loss of VP4 and exposure of negatively-charged amino acid residues in the capsid’s interior after expansion, create a repulsive viral RNA-capsid interface, aiding genome release.

## Results

### Faf-BSA or ions prime intact CVA9 to expanded virions

We studied factors that lead to virion expansion or uncoating by monitoring changes in the viral RNA (vRNA) accessibility to the fluorescent dye SYBR Green II using real-time spectroscopy. SYBR Green II interacts with free non-encapsidated RNA or it can access vRNA due to the pores in the expanded capsids (6, 27). In DPBS, the virions were stable for 3 hours according to the very low fluorescence readout throughout the measurement (Fig. 1A, black solid line; Table 1). Addition of RNase allowed us to distinguish between the signal originating from vRNA released from dissociated virions and the signal from expanded RNA-containing particles where the RNA is protected (Fig. 1A-B dotted lines). For CVA9 in DPBS, the addition of RNase gave lower fluorescence values indicating that either nucleic acids are present in the CVA9 preparation or vRNA is released from a small percentage of the virions and is degraded by the RNase (Fig. 1A, black dotted line). Incubation of CVA9 in 0.01% fatty acid free bovine serum albumin (faf-BSA) induced a fluorescence signal starting at 20 min and reaching its plateau at 40 min (Fig. 1B red solid line; Table 1). Faf-BSA sequesters fatty acids from the virion. Addition of RNase in a parallel measurement shows only slightly decreased fluorescence readout (Fig. 1B red dotted line vs red solid line). This indicates that 0.01% faf-BSA in DPBS efficiently converted CVA9 into expanded particles and only a small fraction of virions dissociated releasing their vRNA (Fig. 1B).

**Table 1.** Final buffer compositions used for treating CVA9 virions.

| Buffer<sup>a</sup> | NaCl | Na<sub>2</sub>HPO<sub>4</sub> | total Na | KCl | KH<sub>2</sub>PO<sub>4</sub> | K<sub>2</sub>HPO<sub>4</sub> | total K | MgCl<sub>2</sub> | CaCl<sub>2</sub> | faf-BSA (μM) | Palmitate (μM) |
| --- | --- | --- | --- | --- | --- | --- | --- | --- | --- | --- | --- |
| PBS-Mg (storage) | 137 | 8 | 153 | 3 | 2 | - | 5 | 2 | - | - |  |
| DPBS | 138 | 8.1 | 154.2 | 2.7 | 1.5 | - | 4.2 | 0.5 | 0.9 | - | - |
| Ion treatment | 20 | - | 20 | - | 6 | 12 | 30 | 0.5 | 0.45 | - | - |
| Ion treatment, 15 μM palmitate | 20 | - | 20 | - | 6 | 12 | 30 | 0.5 | 0.45 | - | 15 |
| 0.01 % faf-BSA<sup>b</sup> | 138 | 8.1 | 154.2 | 2.7 | 1.5 | - | 4.2 | 0.5 | 0.9 | 1.52 | - |
| 0.005 % faf-BSA | 138 | 8.1 | 154.2 | 2.7 | 1.5 | - | 4.2 | 0.5 | 0.9 | 0.75 | - |
| 0.0005 % faf-BSA | 138 | 8.1 | 154.2 | 2.7 | 1.5 | - | 4.2 | 0.5 | 0.9 | 0.075 | - |
| 0.005 % faf-BSA, 15 μM palmitate | 138 | 8.1 | 154.2 | 2.7 | 1.5 | - | 4.2 | 0.5 | 0.9 | 0.75 | 15 |
| 0.01 % faf-BSA in ion buffer (combined) | 20 | - | 20 | - | 6 | 12 | 30 | 0.5 | 0.45 | 1.52 | - |

<sup>a</sup> if not indicated, concentrations are given in mM
<sup>b</sup> 0.01% faf-BSA corresponds to 20-fold excess of faf-BSA molecules per hydrophobic pocket in the virion (60 pockets per virus), when 10 μg/mL of CVA9 is used. This gives approximately 1.25 nM concentration of CVA9 with estimated MW of the virion of 8MDa.

**Figure 1.**
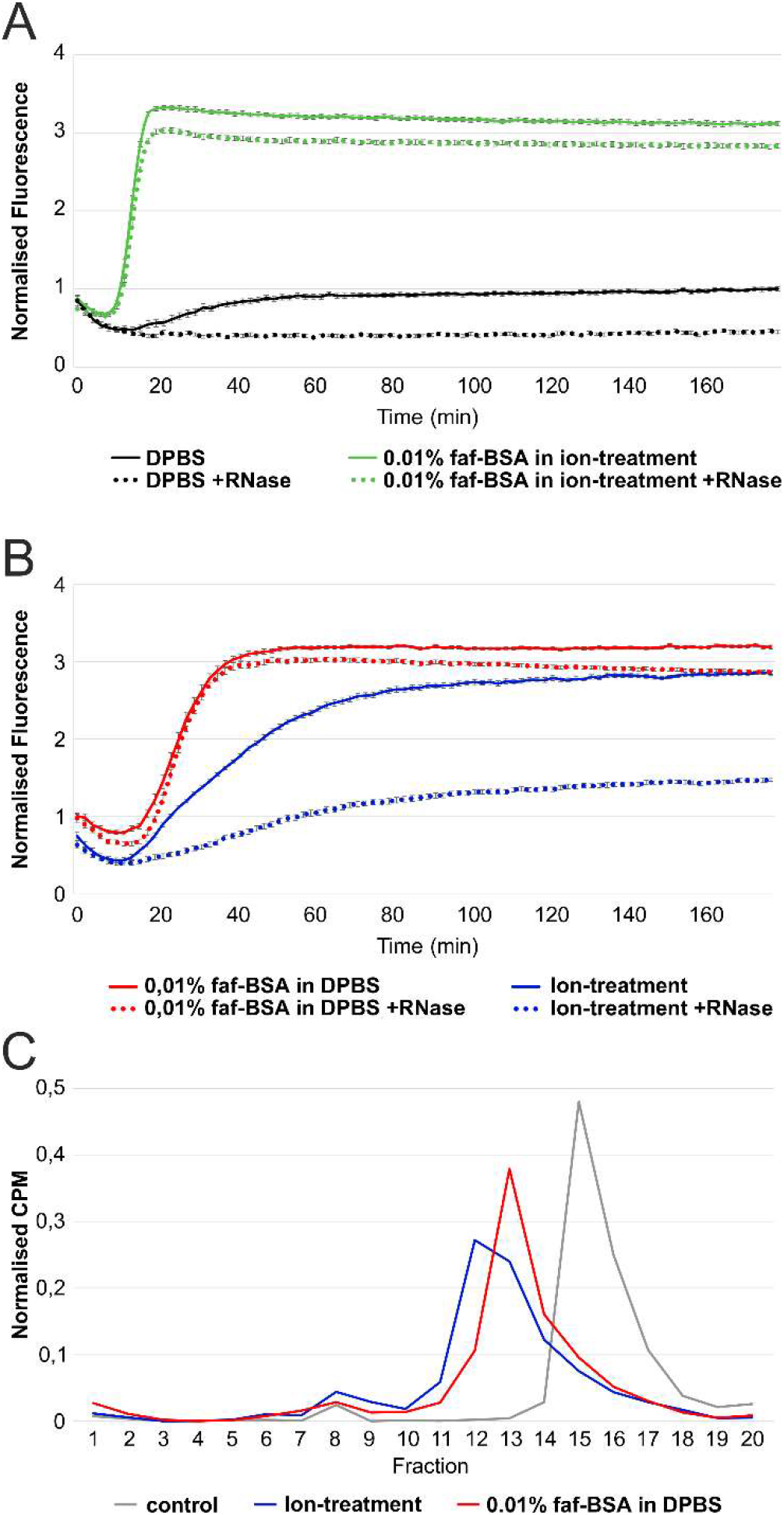
Endosomal ion and 0.01 % faf-BSA treatment, individually or in combination, leads to CVA9 virion expansion. **(A)** Real-time spectroscopy measurements of fluorescent dye SYBR Green II accessibility to CVA9 genome in the presence of 0.01 % faf-BSA in endosomal ionic environment (green) with (dotted line) or without RNase (solid line). Untreated CVA9 in DPBS (black) with (dotted line) or without RNase (solid line) was used as a control. **(B)** Real-time spectroscopy measurements of fluorescent dye SYBR Green II accessibility to CVA9 genome in the presence of 0.01 % faf-BSA (red) or endosomal ion composition (blue) with (dotted line) or without RNase (solid line). The fluorescence signal under the dotted curve indicates expanded virions, whereas fluorescence signal between solid and dotted curves originates from released genome, i.e. empty capsids. **(C)** Effect of 0.01 % faf-BSA and endosomal ions on metabolically labeled ^35^S CVA9 virions as analyzed by ultracentrifugation in 5 % to 20 % sucrose gradient and subsequent fractionation. CVA9 after endosomal ionic treatment (blue) and 0.01 % faf-BSA treatment (red) are shown. Non-treated CVA9 in PBS-Mg buffer was used as a control (grey). Intact viruses are observed mainly in fractions 15 - 16, intermediate particles mainly in fractions 12 - 13 and empty particles in fractions 8 - 10. The exact buffer composition is given in Table 1.

The same method was applied to study the effect of estimated endosomal ionic content on CVA9 virion expansion (Table 1). Ion-treatment caused the fluorescence signal for vRNA to be detected after 15 min, which gradually increased, reaching a plateau in 3 hours (Fig. 1B, blue solid line). The addition of RNase to the ion-treatment reaction reduced the fluorescent signal by half (Fig. 1B blue dotted vs blue solid line) indicating that during ion treatment a significant fraction of the CVA9 virions released their vRNA. Thus, incubation of CVA9 in the buffer mimicking endosomal content induces virion expansion and in addition RNA release from some virions.

The combined treatment caused an additive effect in the fluorescence signal kinetics with expansion plateauing within 15 min (Fig. 1A green lines). However, it considerably reduced RNA release from the virions compared to ion treatment alone (green dotted line in Fig. 1A vs blue dotted line in Fig. 1B).

### Sucrose gradient analysis reveals sample heterogeneity

Sucrose gradient analysis of metabolically-labelled CVA9 confirmed the results of the fluorescence analysis. CVA9 treated with 0.01% faf-BSA in DPBS and separated by differential ultracentrifugation shifted the peak fraction compared to the control (Fig. 1C red vs grey line). This shift indicates the formation of expanded particles as their migration in the sucrose gradient is slower than that of the intact particles (28). In contrast, the ion treatment of CVA9 resulted in a slightly larger shift and a broader peak in the sucrose gradient (Fig. 1C blue line) as well as a larger and broader peak of empty virions (fractions 8-10) in accordance with the fluorescence measurements (Fig. 1B). The overlapping peaks reveal incomplete separation of the particles in the gradient.

### AF4 allows separation of expanded and intact particles

AF4 has been shown to be useful in virus purification instead of ultracentrifugation (29, 30), and in our experiments, AF4 allowed the physical separation of expanded particles (stabilized by faf-BSA) from intact ones and thus biochemical analysis of the proteins present (Fig. 2). Importantly, if VP4 is released from the expanded virions, it will flow through the 10 kDa cut off AF4 membrane used in this experimental setup and will not reach the fraction collector. Thus the protein composition analysis of the virions separated by AF4 will directly reflect only the protein composition in the particle.

**Figure 2.**
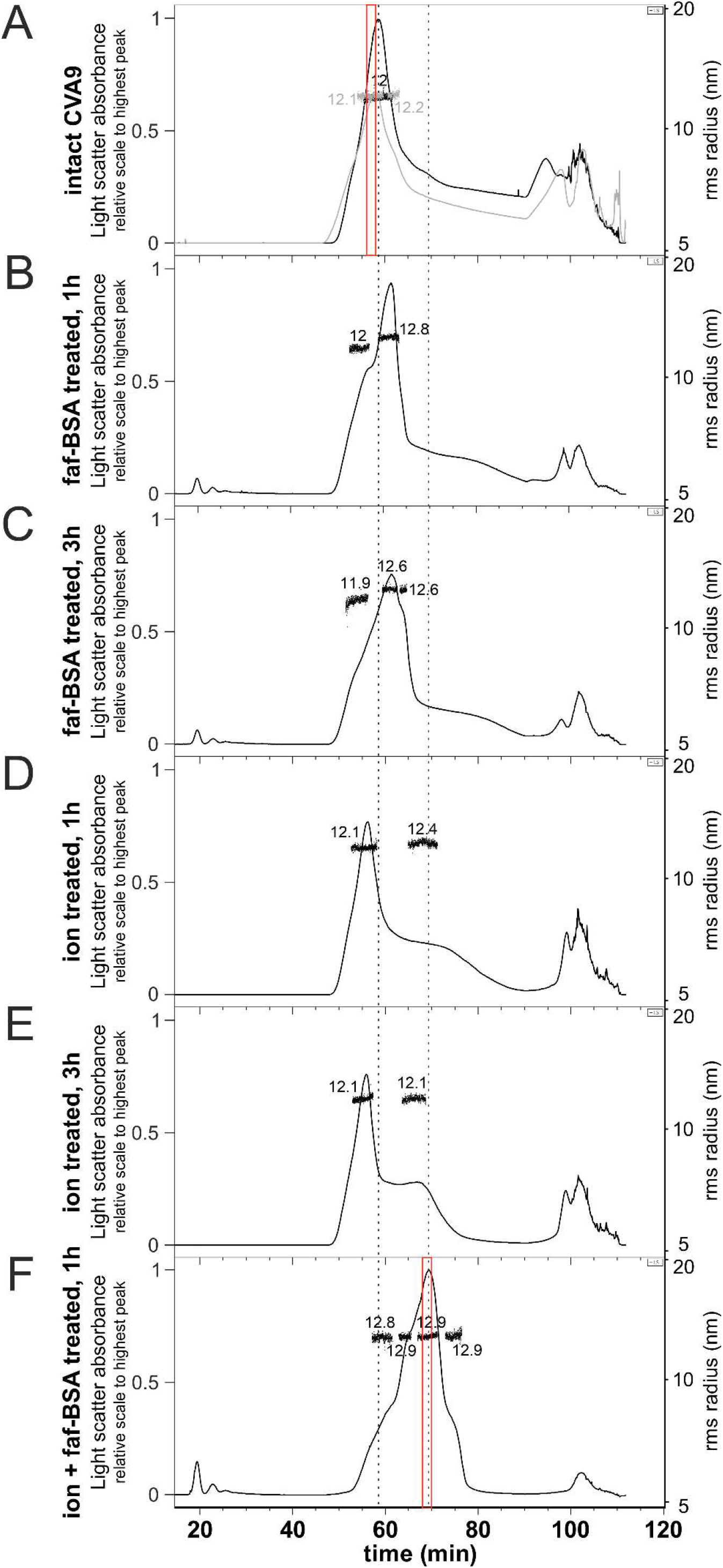
AF4 allows separation of heterogeneous CVA9 virion population generated by 0.01 % faf-BSA and endosomal ion-treatment. **(A)** Two independent AF4 fractograms of untreated CVA9. **(B)** AF4 fractogram of CVA9 after 1 hour treatment with 0.01 % faf-BSA at 37 °C. **(C)** AF4 fractogram of CVA9 after 3 hour treatment with 0.01 % faf-BSA at 37 °C. **(D)** AF4 fractogram of CVA9 after 1 hour treatment in endosomal ion environment at 37 °C. **(E)** AF4 fractogram of CVA9 after 3 hour treatment in endosomal ion environment at 37 °C. **(F)** AF4 fractogram of CVA9 after combined treatment (0.01 % faf-BSA in endosomal ionic environment) for 1 hour at 37 °C. In all panels, x axis indicates time, 1-mL fractions were collected at the rate of 0.5 mL/min; y axis on the left shows light scattering (black or grey lines), and on the right hydrodynamic radius is shown as measured by online dynamic light scattering detector (black dots). Vertical dashed line on the left indicates peak for intact and on the right peak for expanded particles. AF4 fractions used for MS analysis are indicated by red boxes.

The light scattering signal in the AF4 fractograms of the untreated CVA9 showed a clear peak corresponding to particles with a radius of gyration (Rg) of 12 and 12.1 nm (two separate measurements) giving geometric radii (R) of 15.5 and 15.6 nm, respectively (Fig. 2A, Table 2). The tail at 70-90 min elution indicates particle heterogeneity in the sample and the peaks at the end of the fractogram (90-110 min) indicate aggregation. AF4 analysis of 0.01% faf-BSA-treated CVA9 revealed multimodal peaks indicating a few transient states of the virions with lower diffusion coefficients compared to the intact virions (Fig. 2B, C). The Rg of particles corresponding to the highest peak was estimated to 12.8 nm (after 1 h treatment) and 12.6 nm (after 3 h treatment) giving R of 16.5 and 16.3 nm, respectively, indicating expanded virions. The peaks for soluble protein eluting at early time points (about 20 min) correspond to BSA introduced in the buffer for the experiment.

**Table 2.** AF4 Results.

| CVA9 treatment | Rh, nm | Rg, nm | Rg/Rh | R, nm |
| --- | --- | --- | --- | --- |
| no treatment (control) | 14.8 | 12.1 | 0.82 | 15.6 |
| faf-BSA 1h | 17.0 | 12.8 | 0.75 | 16.5 |
| faf-BSA 3h | 16.7 | 12.6 | 0.75 | 16.3 |
| ions 1h | 14.7 | 12.1 | 0.82 | 15.6 |
| ions 3h | 14.7 | 12.1 | 0.82 | 15.6 |
| combined treatment 1h | 17.0 | 12.9 | 0.76 | 16.7 |

Rh, hydrodynamic radius; Rg, radius of gyration; R, geometric radius.
Geometric radius has been estimated using following equation:
$(Rg)^2=3/5R^2$

Similarly to the real time spectroscopy measurements, ion treated-CVA9 showed slow virion conversion to expanded particles, as the main peak corresponds to the intact virions with an estimated Rg 12.1 nm (R of 15.6 nm) and a shoulder at elution time 65-75 min corresponding to expanded virions, for which Rg measurements are unreliable due to the low signal (Fig. 2D, E). The slight shift of the peak for intact virions could be explained by the changes in ion composition introduced by treatment.

Combined treatment with both ion and 0.01% faf-BSA resulted in a clearly shifted peak with a particle population of Rg 12.9 nm (R 16.7 nm) (Fig. 2F). The first shoulder of the peak eluting at 60-63 min (Fig. 2F) could correspond to the main peak of the 0.01% faf-BSA treated CVA9 (Fig. 2B, C). The second shoulder of the sample eluting at about 65 min could correspond to the right shoulder of the CVA9 treated for 3 hours with 0.01% faf-BSA only (Fig. 2F compared with Fig. 2C). In the sample with the combined treatment, as the major peak (53-77 min) has no tail and there is the smallest aggregation peak, we conclude the particles are expanded and stabilized by the treatment. Overall, AF4 enabled us to separate the expanded particles from the intact ones in two distinct peaks, which we then used for determination of the particle’s protein composition by MS.

### Mass spectrometry confirms that most of VP4 is lost in expanded virions

In order to investigate if VP4 is present or absent in the expanded virions, we performed MS analysis on AF4 peak fractions of the untreated CVA9 and CVA9 after combined treatment (indicated in red in Fig. 2A and 2F). The MS results indicate reduced spectral counts for VP4 in the treated vs the control CVA9 when compared with the total counts for the other structural proteins (Suppl. Table 1). In the control sample, 8.8% of the spectral counts for the structural proteins correspond to VP4 (13 spectra out of 134), which agrees well with the stoichiometry-based calculations from the virion amino acid composition where VP4 accounts for 7.8% of the total. In the treated CVA9 sample, the spectra for VP4 reach only 0.9% (5 out of 533) indicating that most of the VP4 is lost in the expanded particles. Due to the fractionation in AF4, free VP4 is removed from the particle peak, thus MS analysis reflects only the protein present in the particles.

### Palmitate prevents particle expansion

The faf-BSA effect on CVA9 particle expansion is concentration dependent. Treating the virus with 0.01% faf-BSA, equivalent to a twenty-fold excess of faf-BSA molecules per hydrophobic pocket, we observed the efficient formation of expanded virions with only a small fraction corresponding to empty capsids as analysed by sucrose gradient differential centrifugation (Fig. 3A, red line vs. grey line; Table 1). A twenty-fold lower concentration of faf-BSA (0.0005%) equal to a 1:1 molar ratio of faf-BSA per hydrophobic pocket, had only a moderate effect on the formation of expanded CVA9 particles (Fig. 3A, orange). Furthermore, in the case of the addition of an excess amount of palmitate (in relation to faf-BSA) to saturate both the faf-BSA and the virions, the CVA9 virions were protected from expansion (Fig. 3B pink). Similar results were obtained when an excess amount of palmitate was added to CVA9 ion-treatment reaction (Fig. 3B blue vs. light blue line).

**Figure 3.**
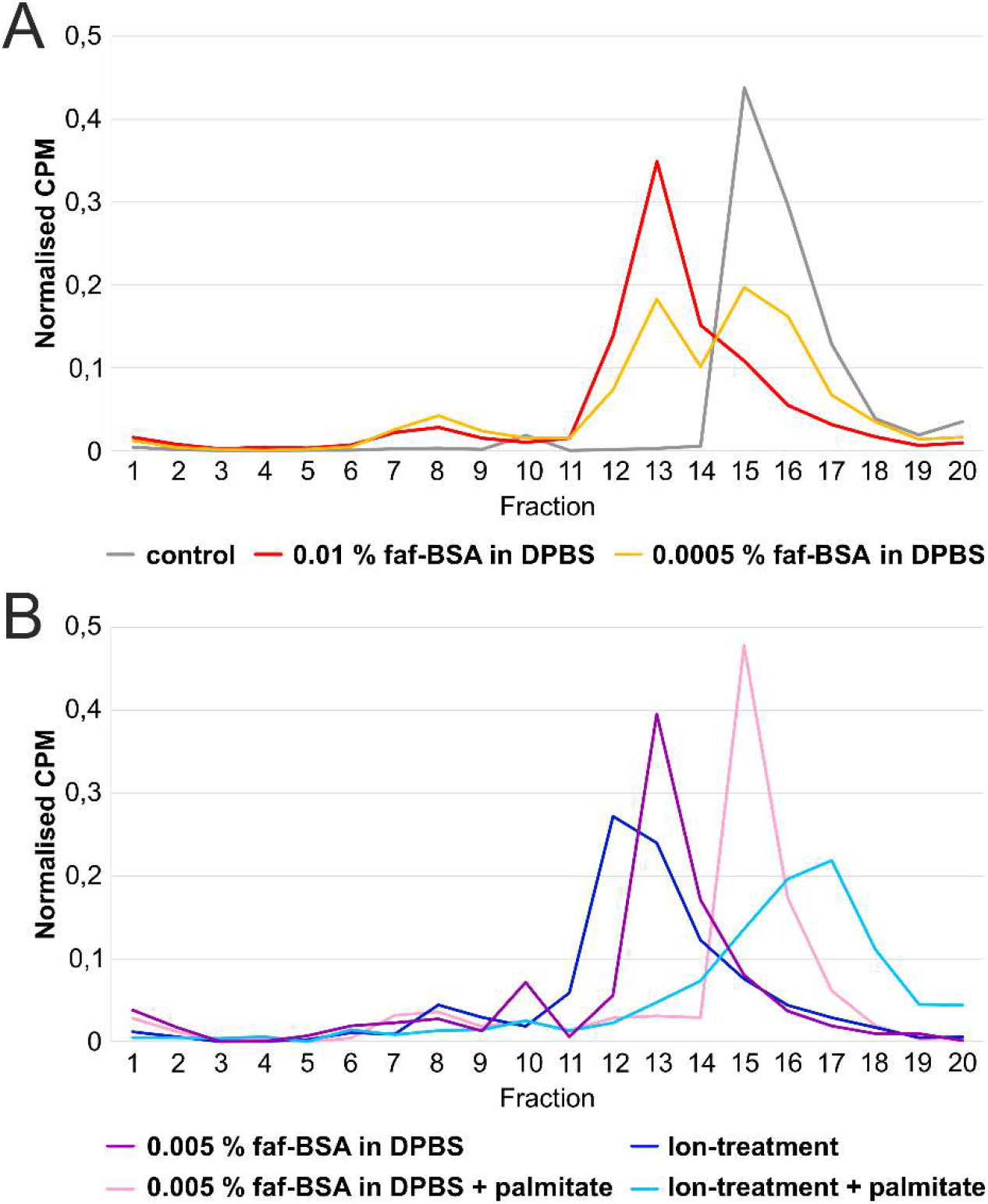
CVA9 expansion dependency on faf-BSA and palmitate concentrations. **(A)** Fractionation of untreated (grey), 0.01 % faf-BSA (red) or 0.0005 % faf-BSA (orange) treated CVA9 after separation in 5 % to 20 % sucrose gradient using ultracentrifugation. Peak for intact virions corresponds to fractions 15 - 17, for expanded virions to fractions 12 - 14, and empty virions to fractions 7 - 10. **(B)** Analysis of CVA9 treated with 0.005 % faf-BSA in the presence of excess of palmitate (pink) after ultracentrifugation in 5 % to 20 % sucrose gradient. For comparison, 0.005 % faf-BSA treated CVA9 ultracentrifuged the same way without addition of palmitate is shown (purple). Similar analysis of CVA9 treated with endosomal ion concentrations with (blue) or without addition of palmitate (light blue) is also shown. Peak for intact virions corresponds to fractions 15 - 18, for expanded virions to fractions 12 - 14, and empty capsids to fractions 7 - 10. Buffer composition is shown in Table 1.

### The heterogeneous particle population can be separated computationally

We used cryo-EM to analyze the intact CVA9 virions which were used as the input for faf-BSA or ion-treatment as well as the endpoint of the treatment achieved after 1 h (for faf-BSA treatment) or 3 h (for ion-treatment) at 37 °C. Multiple rounds of 2D and 3D classifications, including focused classification, revealed several different particle types in all three data sets (Fig. 4 and 5; Table 3). As expected, the main particle population in the control data set was that of intact particles (89%), but there were also expanded particles containing viral genome (5%) and empty non-expanded capsids observed (6%) (Fig. 5A). The low number of expanded and empty particles in the control data set corresponds well with the low fluorescence signal from the untreated CVA9 sample (Fig. 1A black lines); moreover, empty capsids may have released their RNA during storage which was detected in the fluorescent measurement with RNase or there was nucleic acid impurity in the sample (Fig. 1A black dashed line). In faf-BSA and ion-treated CVA9 data sets the main populations, 68% and 94.4% respectively, conform to expanded virions containing the viral genome (Fig. 5B, C). In addition, the faf-BSA-treated CVA9 data set contained a significant number of intact (non-expanded) and empty expanded particles, in agreement with the AF4 results showing a signal for intact particles in addition to expanded ones (Fig. 2B). The ion-treated CVA9 data set had the lowest percentage of intact and empty expanded particles (Fig. 5C). Free RNA detected as a drop in fluorescence signal when RNase was added (Fig. 1B blue dashed line), AF4 results showing a peak primarily for intact particles (Fig. 2D and E), and the overall low particle number picked from the micrographs (Table 3) indicate particle instability after ion treatment. In both, faf-BSA and ion-treated CVA9 data sets, the empty capsids were expanded with an enlarged capsid diameter and openings at the two-folds (Fig. 5B, C).

**Table 3.** Particle heterogeneity.

| type of particles | native CVA9 | BSA-treated CVA9 | ion-treated CVA9 |
| --- | --- | --- | --- |
| total number of particles | 21 365 | 8 061 | 9 055 |
| empty particles | 1 275^ | 853* | 93* |
| expanded particles | 1 024 | 5 533 | 8 540 |
| non-expanded particles | 19 066 | 1 675 | 422 |

\* expanded particles ^ non-expanded particles

**Figure 4:**
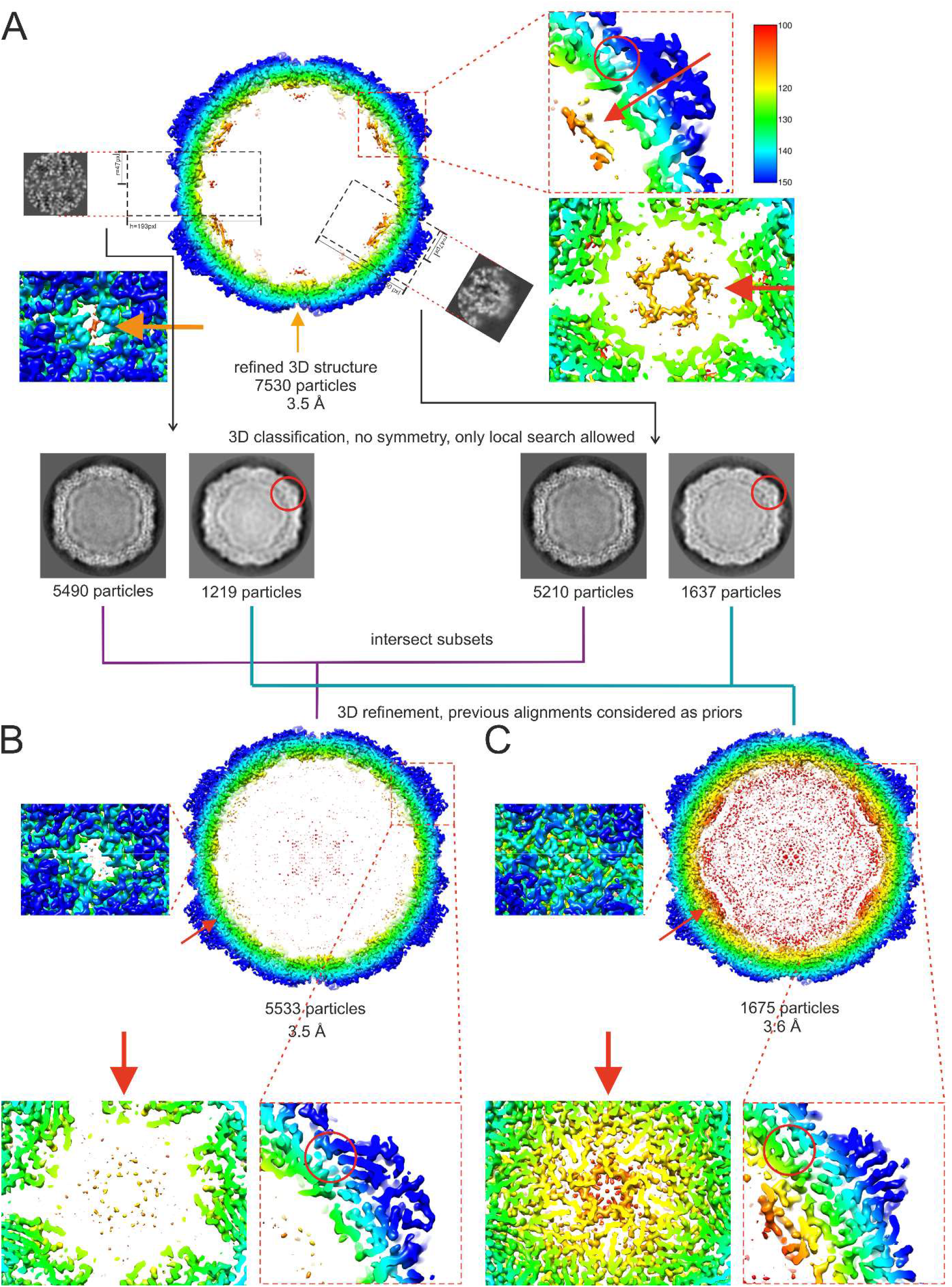
Workflow of focused classification applied to the faf-BSA treated CVA9 dataset to sort heterogeneity. **A**. Cross section of an initial faf-BSA treated CVA9 reconstruction using standard 2D and 3D classifications. Features of interest at 2-fold and 5-fold are enlarged as well as the position of the hydrophobic pocket. Despite the nominal resolution of 3.5 Å resolution, the density under the 5f was much more ordered than expected. Focused classification on either 2-fold or 5-fold yielded two distinct classes corresponding to intact and expanded particles ordered density was due to contamination with virion particles in the reconstruction. The consensus classes were used for further refinement. **B**. Cross section of the faf-BSA expanded CVA9 particle density map. Features at 2-fold, 5-fold and the collapsed hydrophobic pocket are enlarged. **C**. Cross section of the native CVA9 particle density map obtained from a 2D class in faf-BSA treated CVA9 dataset. **A-C**. Features at 2-fold, 5-fold and the collapsed hydrophobic pocket are enlarged. Density maps are radially colored in Å from center according to the color key on the right in **A**. Red circles indicate the position of VP1 where the hydrophobic pocket is occupied in the virion, but collapsed in the expanded particle. Red arrows indicate representations where VP4 is organized in the virion, but missing in the expanded particles.

**Figure 5.**
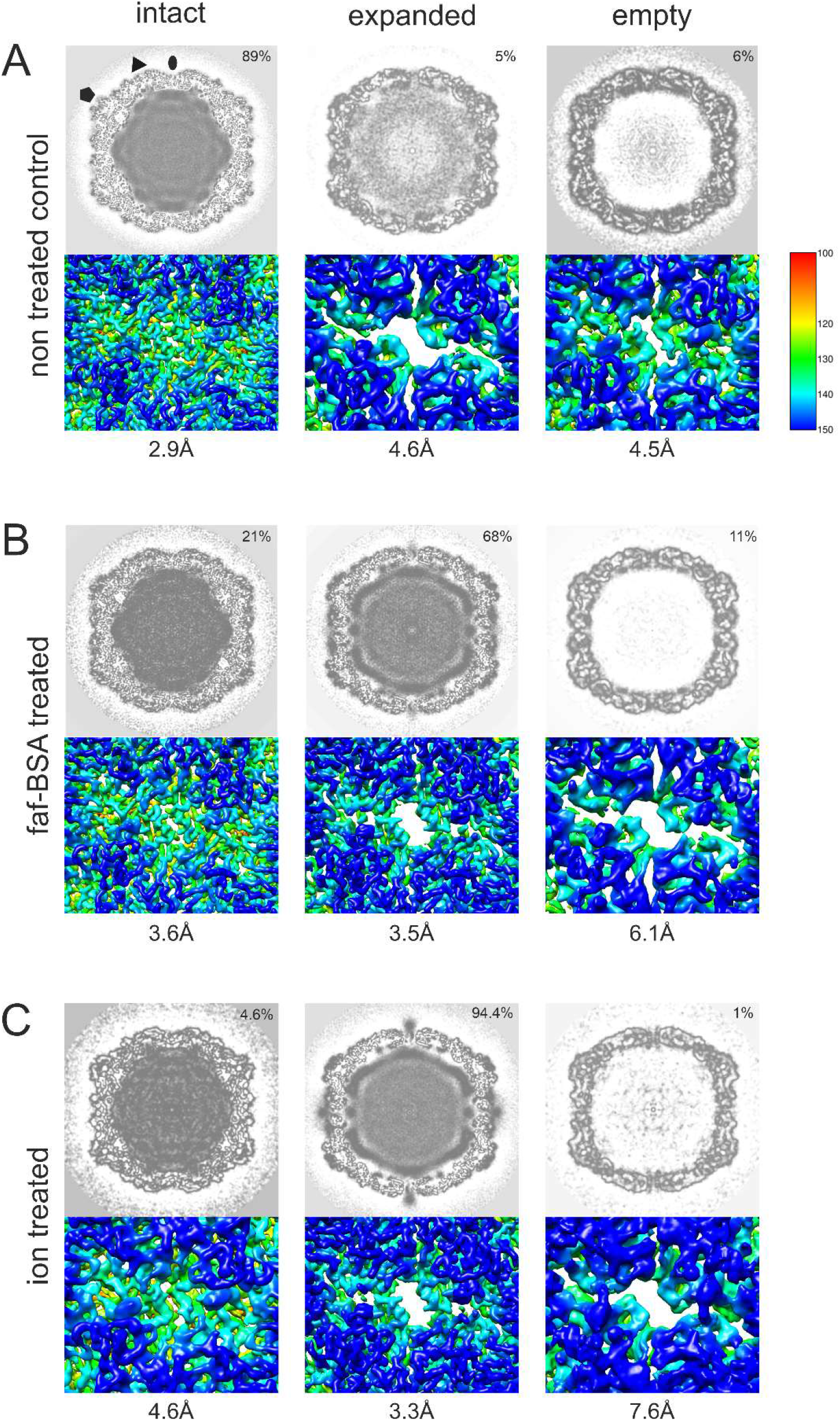
Heterogeneity of CVA9 virions in different data sets after cryo-EM and single particle analysis. **(A)** Analysis of data set of untreated CVA9 resulted in intact, empty non-expanded and expanded virions containing the genome. **(B)** Data set of 0.01 % faf-BSA treated CVA9 separated into expanded empty and filled capsids and in addition to intact particles. **(C)** CVA9 after treatment with endosomal ionic concentrations, in addition to main class of genome-containing expanded virions, yielded small population of intact virions and expanded empty capsids. In all panels, top view shows central plane of the refined reconstructions, and below the enlarged view of a surface representation of a reconstruction is shown at 2-fold axis clearly indicating opening seen in expanded virions. Distribution of particle classes is indicated on the top right corner and resolution of the reconstruction is shown below the figure. Reconstructions are colored according to radial distance in Å from the particle center (color key shown on the left) and shown at 3 σ above mean.

### Cryo-EM shows that faf-BSA or ion treatment of CVA9 leads to A-particle formation

To evaluate structural changes in the CVA9 virions induced by different treatments, we solved cryo-EM structures of the most populated classes from intact, faf-BSA treated and ion-treated CVA9 data sets to 2.9, 3.5 and 3.3 Å resolution, respectively (Table 4; Fig. 6). The well-defined density of the intact CVA9 virion shows features similar to those observed in capsids of other intact enteroviruses and accommodates well the model for all four structural proteins of CVA9 obtained by X-ray crystallography (PDB ID 1D4M) in a T=1 arrangement (29) (Fig. 6 and 7A). Similarly to the X-ray structure, the intact virion reconstruction showed elongated density for a lipid factor placed in the hydrophobic pocket in VP1 (Fig. 7A), as well as a large portion of the myristoyl group covalently attached to the VP4 N-terminus. Furthermore, the base stacking interaction between vRNA and VP2 Trp38 next to the 2-fold axis inside the intact virion could be clearly observed (Fig. 7A). The average radius of the intact virion, 15.6 nm, coincides well with the geometric radius measured in AF4 experiments (Table 2). Compared to the intact virions, the two cryo-EM reconstructions of CVA9 virions treated in two different ways, 0.01 % faf-BSA or ion composition, yielded RNA-containing A-particles expanded by approximately 4% in diameter (Fig. 6 and 7B, C). In addition, these reconstructions revealed more flexibility in capsid proteins and no density for the lipid factor. The MS results indicated that there were only non-stoichiometric amounts of VP4 present and no VP4 could be modelled into the reconstructions (Fig. 7, Table 4). The internal density seen in the difference map was attributed to vRNA (Fig. 7B and C).

**Table 4.** Cryo-EM data collection, refinement and validation statistics.

|  | CVA9 | CVA9-ion treatment | CVA9-BSA treatment |
| --- | --- | --- | --- |
| **Data collection and processing** |  |  |  |
| Magnification | 150000 | 120000 | 150000 |
| Voltage (kV) | 200 | 200 | 200 |
| Electron exposure (e-/Å²) | 30 | 30 | 30 |
| Defocus range settings (µm) | -0.8 – 1.8 | -0.6 – 1.8 | -0.8 – 1.8 |
| Pixel size (Å) | 0.97 | 1.24 | 0.97 |
| Symmetry imposed | I2 | I2 | I2 |
| Micrographs (no.) | 2421 | 1964 | 2472 |
| Initial particle images (no.) | 28818 | 9702 | 15179 |
| Good particles | 21365 | 9055 | 8061 |
| Final particle images (no.) | 19066 | 8540 | 5533 |
| Map resolution (Å) | 2.9 | 3.3 | 3.5 |
| FSC threshold | 0.143 | 0.143 | 0.143 |
| Map resolution range (Å) | 999-1.94 | 999-2.48 | 999-1.94 |
| **Refinement** |  |  |  |
| Map sharpening B factor (Å²) | -50 | -70 | -70 |
| **Model composition** |  |  |  |
| **Non-hydrogen atoms** |  |  |  |
| **Protein residues** |  |  |  |
| VP1 | 1-7, 11-283 | 60-127, 138-203, 206-283 | 54-127, 137-282 |
| VP2 | 10-260 | 14-27, 29-43, 50-259 | 14-27, 30-45, 50-260 |
| VP3 | 1-238 | 3-175, 185-233 | 3-175, 184-233 |
| VP4 | MYR-1-14, 23-68 | - | - |
| Ligands | Lipid factor | - | - |
| **R.m.s. deviations** |  |  |  |
| Bond lengths (Å) | 0.67 | 0.88 | 0.87 |
| Bond angles (°) | 0.67 | 1.00 | 0.99 |
| **Validation** |  |  |  |
| MolProbity score | 1.74 | 1.27 | 1.19 |
| Clashscore | 0.9 | 0 | 0 |
| Poor rotamers (%) | 4.85 | 2.05 | 1.65 |
| **Ramachandran plot** |  |  |  |
| Favored (%) | 89.83 | 90.72 | 90.91 |
| Allowed (%) | 8.33 | 5.86 | 7.18 |
| Disallowed (%) | 1.84 | 3.42 | 1.91 |

**Figure 6.**
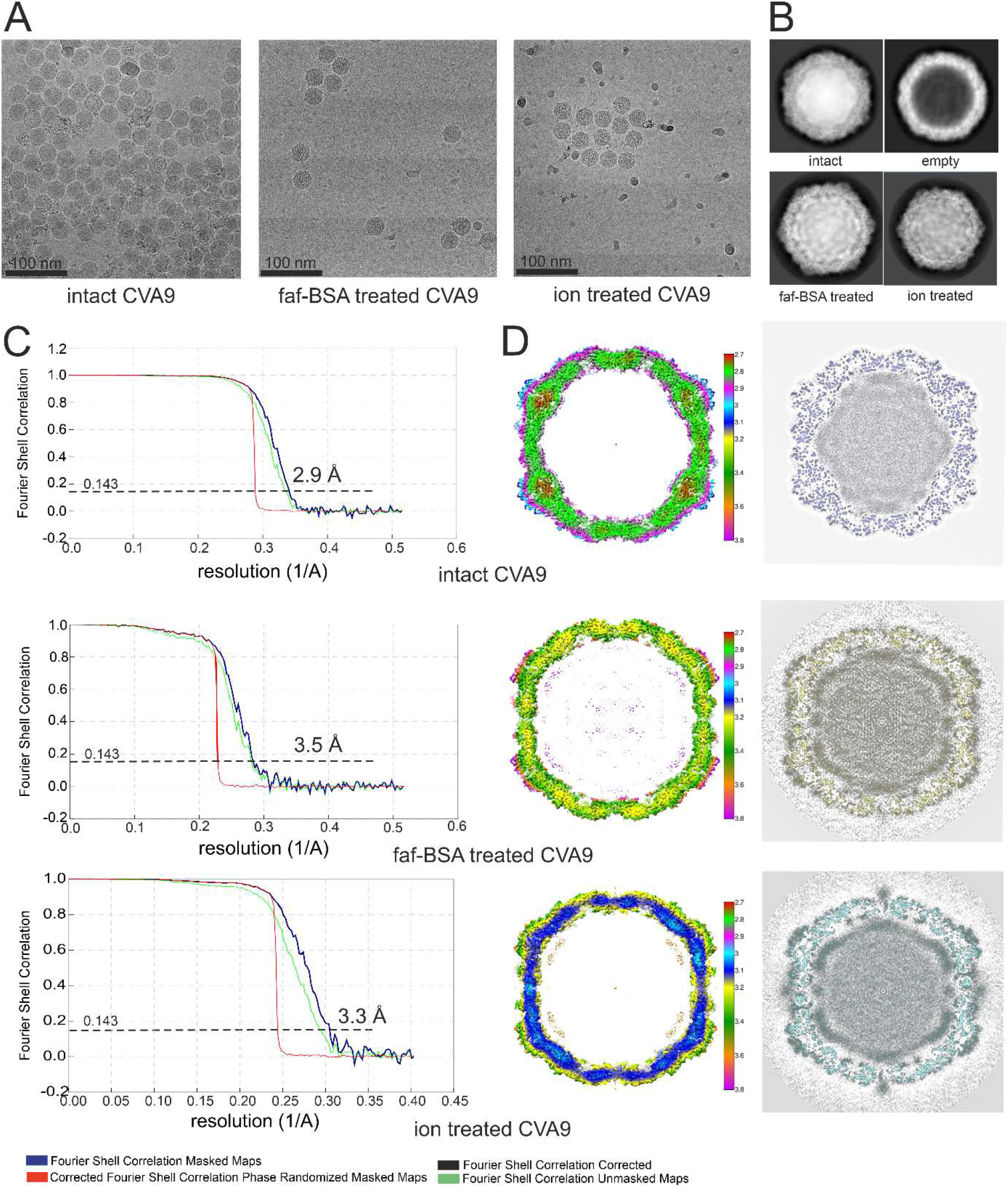
Cryo-EM data collection and single particle analysis. **(A)** Representative micrographs from untreated, 0.01 % faf-BSA and endosomal ionic conditions treated CVA9 data sets. Scale bar, 100 nm. **(B)** Example 2D class averages obtained in Relion 2 during processing workflow for three data sets; 2D class average corresponding to empty virions is from untreated CVA9 data set. **(C)** FSC curves of final reconstructions for intact CVA9, 0.01 % faf-BSA-treated and endosomal ion-treated CVA9 and estimated resolutions at 0.143 threshold. **(D)** Left, central sections of the CVA9 reconstructions colored according to local resolution as shown by the color keys (in Å). Reconstructions are shown at 2 σ above mean. On the right, central planes of the reconstructions are shown.

**Figure 7.**
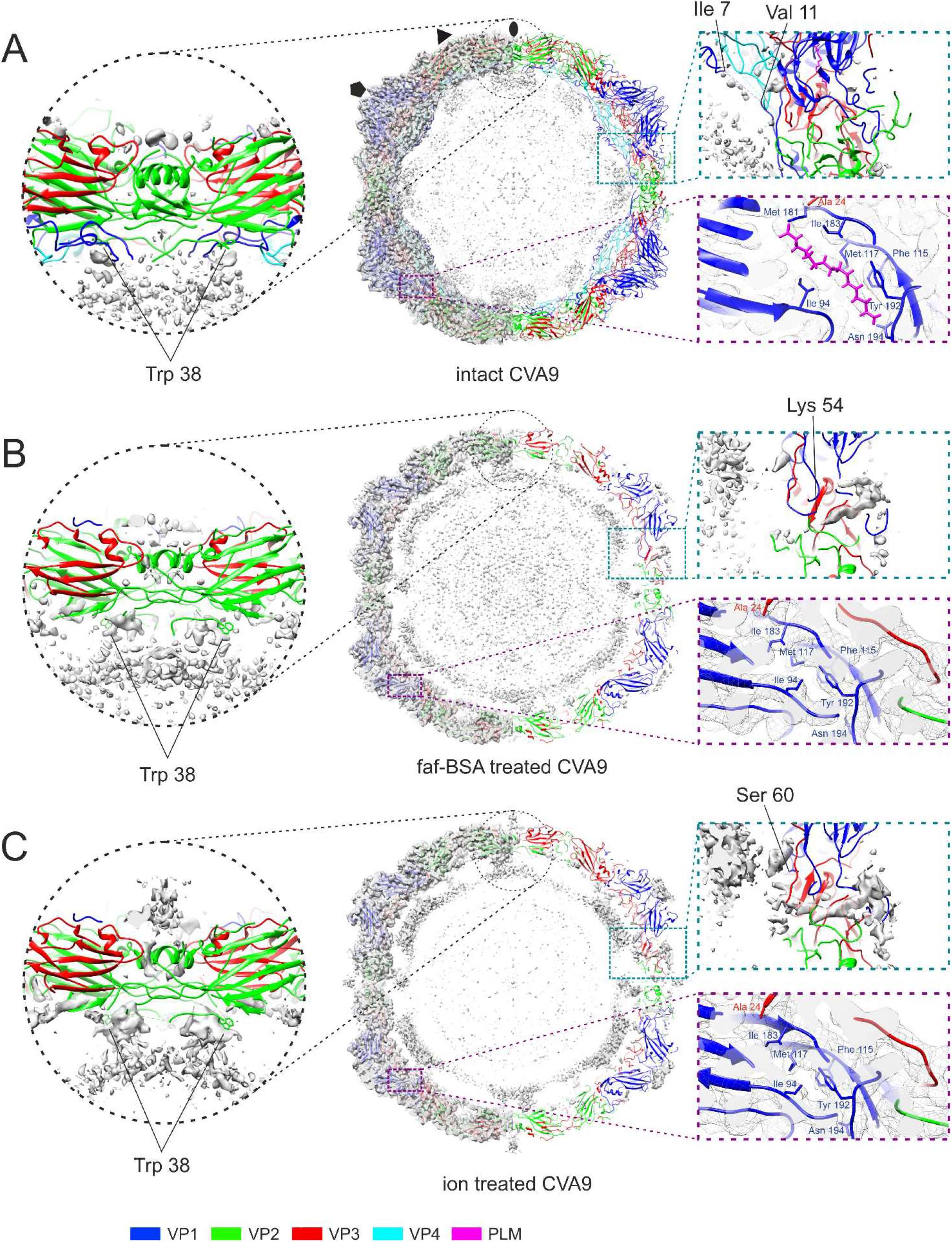
Comparison of reconstructions and atomic models of intact, 0.01 % faf-BSA- and endosomal ion-treated CVA9 virions. **(A)** Untreated CVA9 central section through the model fitted into the reconstruction shown at 3 σ above mean. Left half, cryoEM density with atomic model fit; right half, the cryoEM density accounted for by the atomic model is not shown to reveal the un-modelled density. Symmetry axes are marked as follows, 5-fold (pentagon), 3-fold (triangle) and 2-fold (ellipse). Enlarged view of the model at a 2-fold symmetry axis highlights the vRNA contacts with Trp 38 from VP2 observed in all three reconstructions. Enlarged view on the right shows that most of the density in the capsid region is accounted for. VP1 N-terminus extends on the inside of the capsid hugging VP4 and density for a lipid factor can accommodate an atomic model of a palmitate. **(B)** Central section through the atomic model fitted into the reconstruction of 0.01 % faf-BSA treated CVA9. On the right, half of the density that was accounted by an atomic model was removed to better visualize the un-modelled density. Enlarged view of vRNA contacts with Trp 38 from VP2 is shown. View of un-modelled density near the VP1 N-terminus (Lys 54) spanning the capsid of expanded virions and collapsed hydrophobic pocket are zoomed on the right. **(C)** Central section through the atomic model fitted into the reconstruction of endosomal ion-treated CVA9. On the right, half of the density that was accounted by an atomic model was removed to better visualize the un-modelled density. Enlarged view of vRNA contacts with Trp 38 from VP2 is shown in which the density above 2-fold can also be seen. View of un-modelled density near the VP1 N-terminus (Ser 60) spanning the capsid of expanded virions and collapsed hydrophobic pocket are zoomed on the right.

Further comparison of the two expanded particle reconstructions showed subtle differences between them (Fig. 7B and Fig. 7C). First, less of the VP1 N-terminus is ordered in the ion-treated CVA9 where the first modelled residue is Ser60 compared to Lys54 in faf-BSA-treated CVA9 (Fig. 7). The order of the E1 VP1 N-terminus after combined treatment is similar to that of faf-BSA treated CVA9 (PDB ID: 6O06)(6). In addition, external loops of VP1 are missing Asp128-Asp136 residues in faf-BSA and Asp128-Met137, Gln204, and Arg205 residues in ion-treated CVA9 models (Table 4). In VP2, residues Cys28, Ala29 and Glu46-Ala49 in faf-BSA and Cys28, Asp44-Ala49, and Ala260 in ion-treated CVA9 models are missing (Table 4). In VP3, surface exposed residues Tyr176-Glu183 in faf-BSA and Tyr176-Tyr184 in ion-treated CVA9 models are missing (Table 4). Second, a poorly defined density is seen on the ion-treated, but not BSA-treated, surface above the 2-fold opening (Fig. 7B, C). The 3D classification of particles from ion-treated CVA9 refinement focused on the 2-fold symmetry axis did not result in a better-defined density above the 2-fold opening, preventing a detailed atomic interpretation (Fig. 4). Altogether cryo-EM reveals more flexibility in the ion-treated compared to the faf-BSA-treated CVA9 capsid. This is in agreement with the fluorescence measurements showing that ion treatment of CVA9 virions leads to more unstable A-particles, which lose their vRNA becoming accessible to RNase (Fig. 1B blue lines). In contrast, the faf-BSA treatment of CVA9 generates more stable A-particles as seen by fluorescence assay and AF4 (Fig. 1 and 2).

The average distribution of vRNA in the capsid changes with expansion (Fig. 6). In the intact virion, the RNA is tethered to the capsid via the VP2 Trp38, but also is in close contact all over the capsid interior. Following faf-BSA or ion-treatment, the average shape of the RNA expands, reflecting the shape of the expanded capsid. However, apart from the tethering at the Trp38, the intimate contact is lost (Fig 6). Electrostatic potential calculations for the capsid proteins in intact and expanded virions indicate a significant shift to a much more negatively-charged surface inside the expanded capsid generating repulsive vRNA-capsid interactions that could account for the observed changes and contribute to vRNA exit (Fig. 8).

**Figure 8.**
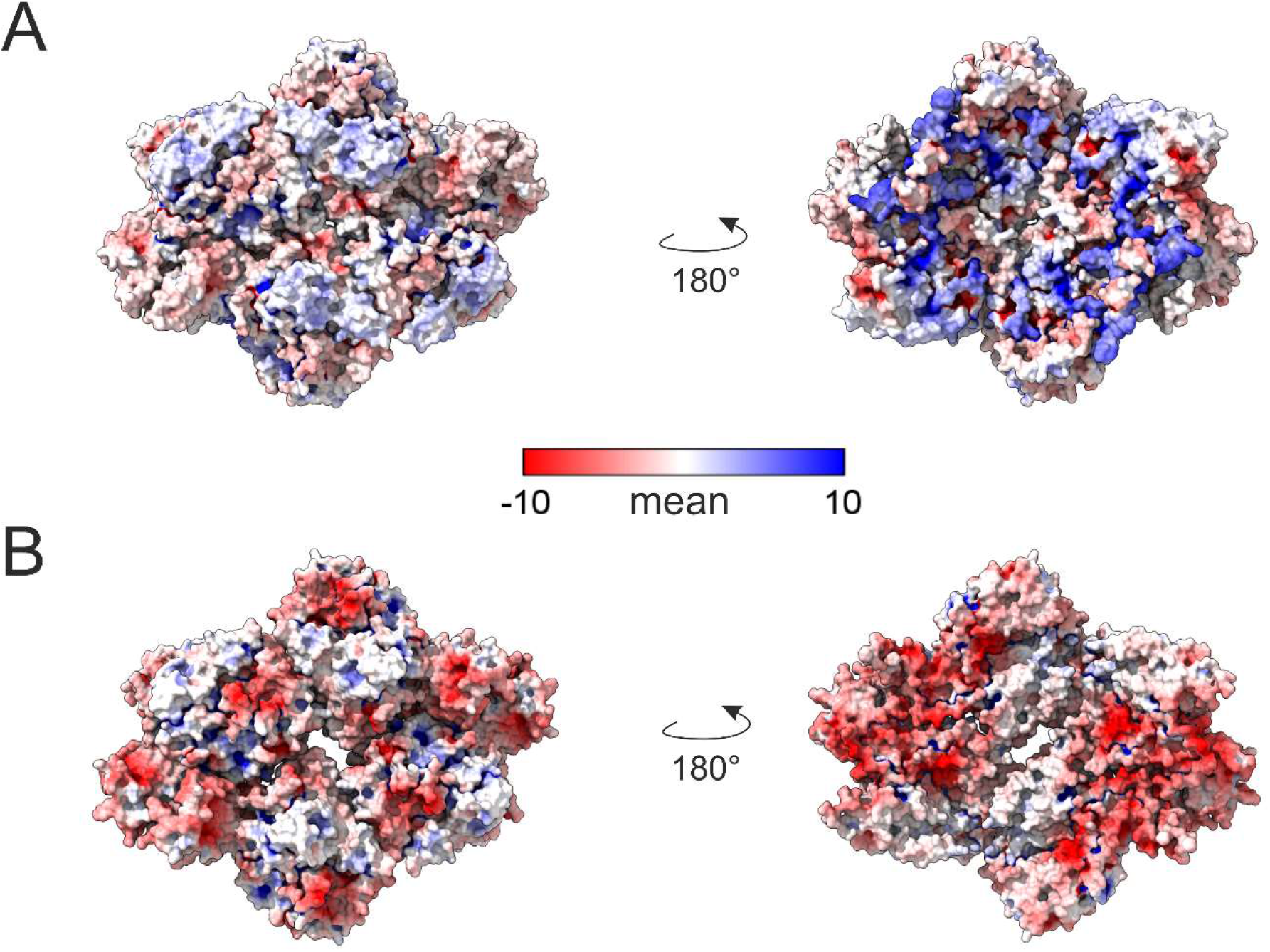
Calculations of electrostatic potentials show significant change from a positive to a negative charge at the inner surface of the capsid when intact and expanded virions are compared. Electrostatic potentials calculated for an asymmetric unit shown for outer (left) and inner (right) surface of untreated **(A)** and 0.01 % faf-BSA treated CVA9 **(B)** as a colored surface according to a color key. Four subunits around 2-fold axis are shown.

## Discussion

In enteroviruses E1, E12 and coxsackievirus B3 serum albumin promotes virion expansion in a dose-dependent manner by sequestering the lipid factor from the VP1 hydrophobic pocket as shown in *in vitro* experiments (6, 20, 21). Here we studied and compared the roles of serum albumin and endosomal ion concentrations at neutral pH in triggering CVA9 expansion and genome release. We show that when used independently, both albumin and endosomal ion concentrations cause particle expansion. However, the rate and extent of the order depend on the condition. The CVA9 treatment with albumin induces expansion in the majority of the particles exposing the VP1 N-termini in about 30 min where most of the particles still contain vRNA (Fig. 1B, Fig. 2, Fig. 7B). The ion treatment leads to a relatively slower particle expansion driving the capsid to a more disordered state and the loss of vRNA from a significant population of particles (Fig. 1B, Fig 2, Fig. 7C, Table 4). Both effects can be prevented by an excess of palmitate preventing the collapse of the VP1 hydrophobic pocket, a necessary event for expansion (Fig. 3B). When virions are treated with faf-BSA, the lipid factor is bound more strongly by the albumin than by the capsid, causing the capsid to expand. In the case of ion treatment, lipid factor dissociation from the particle might be driven by changes in counter ions so that the vRNA swells, destabilizing the capsid for expansion.

A number of pieces of evidence indicate that the endosomes are the final location for picornavirus uncoating and vRNA release into the cytosol (30–33). Our experiments support this idea by showing that changes in cation concentrations to those mimicking the endosomal environment slowly convert CVA9 into expanded virions. Moreover under these conditions, a significant fraction of the virions lose their vRNA or dissociate completely as shown by AF4 and real-time fluorescence measurements (Fig. 1 and 2). Viral RNA genomes inside the capsids are extremely compact despite the repulsive negative charges of the RNA backbone phosphates (34). Tight vRNA folding depends on both the RNA sequence specificity and the presence of cations, which compensate for the repulsive interactions between negatively charged RNA backbones enabling close juxtaposition of two phosphate groups. RNA phosphate oxyanions have significant affinity to Mg^2+^ due to its size and charge density which allows small interatomic distances in RNA-Mg^2+^ clusters giving high stability (35). RNA association with Na^+^ and Ca^2+^ is not as stable, because of the smaller charge of Na^+^ and the bigger size of Ca^2+^ ions. The most abundant cations that associate with highly folded RNAs are Mg^2+^ and Na^+^ (36). As mentioned before, during endosome maturation there is a tendency for decrease in Na^+^ and Ca^2+^ ions and increase in K^+^ ions (22, 23). There is no information about changes in Mg^2+^ concentration in endosomes. Because of the changes in ion composition in endosomes, K^+^ ions might replace Na^+^ and perhaps Mg^2+^ ions in the compact vRNA leading to conformational changes and expansion or swelling of the vRNA. Furthermore, the comparison of electrostatic potential on the internal surface of intact and expanded virions shows that vRNA-capsid interactions become repulsive in expanded virions due to conformational changes and the loss of much of the VP4, thus further fostering genome release (Fig. 8). In support of this, altered vRNA-capsid interactions in an A-particle of human rhinovirus 2 have been described (37).

In conclusion, the enterovirus capsid expansion required for release of the genome in the endosome can be triggered by albumin in the serum, but equally it can be triggered in a receptor- and acidic pH-independent manner by the environment of the endosomal lumen.

## Materials and methods

### Virus production and purification

Green monkey kidney cells (GMK) were purchased from the American Type Culture Collection (ATCC) and were maintained in Eagle’s minimum essential medium (MEM) supplemented with 10 % fetal bovine serum (FBS), 1 % GlutaMAX, and 1 % antibiotic–antimycotic solution in a chamber environment adjusted to 37 °C and 5 % CO_2_. For CVA9 production semi-confluent 5-layer bottles of GMK cells were infected with CVA9 (ATCC, Griggs strain GenBank: D00627.1) using an MOI of 3.8 in medium containing 1 % FBS until total cell detachment (at 37 °C with 5 % CO_2_ for 20 - 24 h). The cells and medium were collected, freeze-thawed three times and the cellular debris was centrifuged into a pellet (JA-10 rotor, 6080 rpm, 30 min, 4 °C). The virus containing supernatant was mixed with polyethylene glycol 6000 (Sigma Aldrich, Saint Louis, Missouri, US) (8 % wt/v) and NaCl (2.2% wt/v) at 4 °C overnight. The precipitate was pelleted (JA-10 rotor, 8000 rpm, 45 min, 4 °C) and resuspended in R-buffer (10 mM Tris-HCl (pH 7.5), 200 mM MgCl_2_, 10 % (wt/v) glycerol). The resuspension was treated with 0.3 % (wt/v) sodium deoxylate (Sigma Aldrich) and 0.6 v/v Nonidet P-40 (Sigma Aldrich) for 30 min on ice and centrifuged (TX-200 rotor, 4700 rpm, 15 min, 4 °C) to pellet the remaining cellular membranes. The virus containing supernatant was loaded on top of linear 10 mL 5-20 % sucrose gradient prepared in R-buffer and centrifuged (SW-41 rotor, 35000 rpm, 2 h, 4 °C). The gradient was collected into 500 μL fractions and the virus containing fractions were detected using NanoDrop 1000 Spectrophotometer (Thermo Scientific, Waltham, Massachusetts, US). The virus containing sucrose fractions were diluted at least 1:5 in 2 mM MgCl_2_-PBS and the virus was pelleted by centrifugation (70Ti rotor, 35000 rpm, 2 h, 4 °C) and resuspended in 2 mM MgCl_2_-PBS. The concentration of the purified virus sample was measured using NanoDrop and the stock was stored at −80 °C in small aliquots.

### ^35^S labelled virus production and purification

The 35S labelled virus was produced in semi-confluent 80 cm^2^ cell culture bottles. The bottles were infected with CVA9 (ATCC, Griggs strain) in low-methionine-cysteine medium supplemented with 1 % FBS for three hours, after which the culture medium was changed into a low-methionine-cysteine medium supplemented with 1 % FBS containing 50 μCi/mL [^35^S] methionine-cysteine. After total cell detachment (20 - 24 h) the cells were collected and freeze-thawed three times. After pelleting the cellular residue (TX-200 rotor, 4000 rpm, 15 min, 4 °C) the supernatant was collected and Tween was added in a final concentration of 0.1 %. Solution was incubated on ice for 30 min, centrifuged (TX-200, 4700 rpm, 15min, 4 °C) and loaded on top of a 2 mL 40 % sucrose cushion on ice. The cushion with the virus was centrifuged (SW-41, 30,000 rpm, 2.5 h, 4 °C), after which liquid above the cushion and 500 μL fraction from the cushion was collected to waste and the remaining three virus containing 500 μL fractions were diluted into 2 mM MgCl_2_-PBS and the virus was pelleted (70Ti, 35000 rpm, 3 h, 4 °C). The pellet was resuspended into 1.5 mL of 2mM MgCl_2_-PBS and separated in 5 - 20 % sucrose gradient in R-buffer (SW41, 35000 rpm, 2 h, 4 °C). After collecting 500 μL fractions, the virus was identified using liquid scintillation analyzer TRI-carb 2910 TR (Perkin Elmer, Waltham, Massachusetts, US). The virus containing fractions were collected and pelleted as earlier. The pellet was dissolved in 2 mM PBS-MgCl_2_ and stored at −80 °C in small aliquots.

### Analysis of treated CVA9 by ultracentrifugation in sucrose gradient

1 μg of un-labelled purified CVA9 was mixed with 1 - 3 μL of ^35^S labelled CVA9 (showing approximately 1000 CPM) and faf-BSA (Sigma-Aldrich) or buffer giving final concentrations as indicated in Table 1 was added to obtain final volume of 100 μL. For molar ratio calculations between the pocket, faf-BSA and palmitate (Sigma-Aldrich), an estimated molecular weight of 8 MDa was used for CVA9 and the ones provided by the supplier for faf-BSA and palmitate. Samples were incubated at 37 °C for 1 h (faf-BSA treatment) or 3 h (ion-treatment) before gradient separation. The control samples (purified CVA9 in storage buffer, Table 1) were kept on ice during the incubations. After the incubation the samples were pipetted on top of linear 5 - 20 % sucrose gradients prepared in R-buffer and centrifuged (SW41, 35000 rpm, 2 h, 4 °C). After centrifugation, 500 μL fractions were collected and mixed with scintillation cocktail (Ultima Gold MW, Perkin Elmer) and the radioactivity of each fraction was measured (Tri-Carb 2910 TR, Perkin Elmer).

### AF4

The AF4 experiments were performed using the Eclipse NEON FFF system (Wyatt Technology, Santa Barbara, USA) composed of Agilent 1260 Infinity II pump and autosampler, analytical long channel with Dilution Control ModuleTM (Wyatt Technology, Santa Barbara, USA), and Agilent 1260 fraction collector. AF4 device was coupled to Optilab^®^ refractive index (dRI) detector (Wyatt Technology, Santa Barbara, USA), DAWN multiangle light scattering detector (MALS) with 18 angles (Wyatt Technology, Santa Barbara, USA), online dynamic light scattering detector (DLS) embedded in one of the DAWN angles (Wyatt Technology, Santa Barbara, USA), and UV detector (Agilent) used at 280 nm. Regenerated cellulose membrane at 10 kDa cut off (Wyatt Technology, Santa Barbara, USA lot RIJB19432) and Technology 525 μm spacer (Wyatt Technology, Santa Barbara, USA lot 247096) was used as the accumulation wall. Focusing was performed at cross flow velocity of 2 mL/min. Sample was injected at 0.2 mL/min. Elution was performed using constant cross flow velocity of 3 mL/min for 80 min; channel flow was 1 mL/min; detector flow was 0.5 mL/min. The AF4 flows were controlled via VISION 3.1 software (Wyatt Technology, Santa Barbara, USA). Channel temperature was set to 25 °C. DPBS filtered through 0.1 μm filter was used as a mobile phase for all experiments. Before sample runs, system set up was validated by running 30 μg of BSA solution (2 mg/mL). For all experiments, sample corresponding to 10 μg of purified CVA9 was injected. ASTRA 8.1 software was used for data analysis.

For AF4 analysis CVA9 was treated essentially the same way as for cryo-EM sample preparation. Briefly, 12 μg of purified CVA9 (0.8 μg/μL) were mixed with faf-BSA at a final concentration of 0.01 % in DPBS giving 136 μL total volume. The sample was incubated either for 1 or 3 hours at 37 °C prior to injecting to AF4 device. To analyse the effect of endosomal ionic content, purified CVA9 virions (12 μg) were incubated in ion-treatment buffer (136 μL total volume) for either 1 or 3 hours at 37 °C and then analysed by AF4. Combined treatment was done by mixing purified CVA9 (12 μg) with 0.01 % faf-BSA in ion-treatment buffer in 168 μL total volume and incubated for 1 hour at 37 °C, prior to AF4 analysis in 20 μL (for control), 140 μL (for ion-treated and combined treatment), or 113 μL (faf-BSA treated) volume.

Results obtained by multi-angle light scattering (MALS) and light scattering detectors were used to calculate particle radius of gyration (Rg) and hydrodynamic radius (Rh), respectively. The Rg/Rh ratio is between 0.75 and 0.82 indicating that shape of the particles present in different samples corresponds to a hard sphere confirming expected particle shape in the peak fractions (Table 2) (38, 39).

### Real-time fluorescence measurements

Conversion of CVA9 virions to expanded and empty or dissociated particles, which release their RNA, was observed by real-time spectroscopy as described earlier (6, 27). In short, 1 μg of virus was mixed with buffers containing different ion and supplement (faf-BSA, palmitate) concentrations in a presence of 10X SYBR Green with or without RNase A. Buffer details are indicated in Table 1. The total volume of each reaction mixture was 100 μL. The increase in fluorescence was observed using PerkinElmer 2030 Multilabel Reader Victor X4 (Perkin Elmer) with F485 lamp filter and F535 emission filter for three hours at 37 °C, and the fluorescence with and without RNase was compared to estimate whether the virus was in intact, expanded or dissociated form. A blank measurement with all factors other than the virus was registered and subtracted from the corresponding values obtained during the measurement with virus containing samples. The obtained fluorescence values were normalised against the DPBS measurement and are shown in arbitrary units (AU).

### Liquid chromatography-mass spectrometry (LC-MS) analysis

The peak fraction corresponding to the intact virions in AF4 run of untreated CVA9 (Fig. 2A) and the peak fraction corresponding to the expanded virions in AF4 run of CVA9 after combined 0.01 % faf-BSA and ion-treatment (Fig. 2F) were used for LC-MS analysis. First, each fraction (1 mL in total) was heated at 60 °C for 5 min to inactivate the virus. Then 1 mL of lysis buffer (8 M Urea buffer, 50 mM NH4HCO3, phosphatase inhibitors (Sigma-Aldrich, P2745), and protease inhibitors cocktail (Sigma-Aldrich, P8340)) was added to each fraction on ice. Undissolved particles were removed by centrifugation at 16,000 × g for 10 min at 4 °C. Samples were reduced with Tris(2-carboxyethyl)phosphine (TCEP; Sigma Aldrich), and alkylated with iodoacetamide. Samples were diluted with 50 mM ammonium bicarbonate (AMBIC; Sigma-Aldrich, 213-911-5) to reduce the urea concentration to < 2 M. Sequencing Grade Modified Trypsin (Promega, PRV5111) was added to the samples and incubated for 16 hours at 37 °C. Finally, the trypsin digested samples were desalted with C18 BioPureSPN Mini columns (Nest Group, HUM S18V). A detailed description of the method used here was previously published (40).

The desalted samples were analyzed using an EvoSep One liquid chromatography system coupled with a hybrid trapped ion mobility quadrupole TOF mass spectrometer (Bruker timsTOF Pro2). The samples were directly loaded on Evotips and separated with an 8 cm × 150 μm analytical column (EV1109, Evosep) using the 60 samples per day method (21 min gradient time). Mobile phases A and B were 0.1 % formic acid in water and 0.1 % formic acid in acetonitrile, respectively. The MS analysis was performed in the positive-ion mode using data-dependent acquisition (DDA) in online parallel accumulation-serial fragmentation (PASEF) mode (41) with 10 PASEF scans per topN acquisition cycle. Raw data (.d) from timsTOF Pro were searched using MSFragger (42) against the home-made CVA9 entries of the capsid proteins. Sequence of CVA9 Griggs isolate was used (43). Carbamidomethylation of cysteine residues was used as static modification. Aminoterminal acetylation and oxidation of methionine were used as the dynamic modification. Trypsin was selected as enzyme, and maximum of two missed cleavages were allowed. Both instrument and label-free quantification parameters were left to default settings.

### Cryo-EM sample preparation and data collection

To analyse the effect of faf-BSA on CVA9 virion structure, 3 μg of purified CVA9 (0.8 μg/μL) were mixed with faf-BSA at a final concentration of 0.01 % in DPBS giving 34 μL total volume. The sample was incubated for 1 hour at 37 °C prior to plunging. To analyse the effect of endosomal ionic content on CVA9 virion structure, purified CVA9 virions (3 μg) were incubated in ion-treatment buffer (34 μL total volume) for 3 hours at 37 °C. For cryo-EM sample preparation, 3.0 μL of above-described samples or non-treated CVA9 virions as a control were applied to glow-discharged grids (Quantifoil R1.2/1.3 holey carbon on 300 copper mesh) that were manually blotted with filter paper and flash-frozen in liquid ethane using a homemade plunger. Data was acquired at the Biocenter Finland National Cryo-EM facility with a FEI TALOS Artica transmission electron microscope operated at 200 kV. Movies were collected with a Falcon III direct electron detector at a nominal magnification of 150 000x (for CVA9 control and CVA9 treated with faf-BSA) or 120 000x (for CVA9 treated with endosomal ionic composition) giving a pixel size of 0.97 or 1.24 Å per pixel, respectively. Each movie consisted of 30 frames with an exposure time of 1 second per frame. The total electron dose was approximately 30 electrons per Å^2^. Movies were corrected for beam-induced motion by aligning and averaging all frames with Motioncor2 implemented in Scipion version 2.0 (44, 45).

### Image processing

The contrast transfer function of the averaged micrographs was estimated by CTFFind4 in Scipion version 2.0 (46, 47). Micrographs with astigmatism and diffraction were excluded from further processing. For particle picking 2-step Xmipp3 was used from Scipion version 1.2, followed by 2D classification in Relion 2 (48–50). Initial 3D models were generated using 2000, 1005 and 8012 particles for CVA9 control, CVA9 ion expanded and CVA9 BSA expanded, respectively, utilising 3D initial model protocol in Relion 2 (49, 51, 52). Particles from the best 3D classes were further refined and post-processed in Relion 2. After post-processing, particles were subjected to CTF-refinement and Bayesian polishing using default parameters in Relion 3 (49, 51–54). Local resolution of the maps was calculated in Relion 3 at 15 Å sampling rate and 25 Å resolution threshold for randomizing phases (53, 55). However, at this stage, inspection of a 7530 particle reconstruction of the BSA-treated CVA9 data set showed strong but poorly defined density beneath the capsid near the 5-fold axis (Fig. 4). To try to resolve this 5f density we performed focused classification at the 5-fold symmetry axis (Fig. 4). For this a cylindrical mask of 46 Å radius and 145 Å height was applied with a centre shift below the 5-fold symmetry axis and only a small angular search was allowed (48, 56). The focused 3D classification separated all particles into two well-defined classes. Classes were further refined and post-processed in Relion 3 resolving characteristic features of expanded or intact particles while the density below the 5-fold seen in the initial faf-BSA-treated CVA9 reconstruction disappeared and was attributed to VP4 density from contaminating intact particles (Fig. 4). Approximately 18 % of all particles in the initial BSA-treated data set were found to be non-expanded intact virions (Table 3). Similar results were obtained by 3D classification of the same set of particles focused on a 2-fold symmetry axis by applying cylinder mask of 45 Å radius and 187 Å height and shifting the centre of the mask so that capsid region at the 2-fold axis would be covered (Fig. 4). A similar workflow of focused 3D classification was also applied to ion-treated CVA9 and control data sets, but did not result in any other distinct classes other than the major class. Although this procedure seemed to be robust in separating the intact and expanded particles, it was still not sufficient. So additional rounds of inspection in UCSF Chimera, and classification in both 2D and 3D were carried out, culminating in the identification of intact virions, non-expanded empty particles, expanded virions and expanded empty particles. The majority particle type for the control and the two treatments were then CTF refined and post-processed in Relion 3.

The FSC curve was calculated and the resolution of the final reconstructions was determined according to the gold-standard FSC = 0.143 threshold criterion (57). The image processing and refinement statistics for each data set are shown in Table 4 and the focused classification workflow is shown in Fig 4.

### Modelling and analysis

The CVA9 atomic model PDB ID 1D4M was fitted into the CVA9 control reconstruction using UCSF Chimera 1.15 and manually adjusted in Coot 0.9.2 (58, 59). The initial model for the CVA9 A-particle was a homology model calculated in I-Tasser using the expanded E1 particle PDB ID 6O06 as a reference (6, 60). I-Tasser derived models were fitted into the density maps of faf-BSA treated and ion treated CVA9 using UCSF Chimera 1.15 and were manually adjusted in Coot 0.9.2. Residues for which densities were not observed in the cryo-EM maps, were deleted as indicated in Table 4. All three optimized models were further refined against the cryo-EM map in Molecular Dynamics Flexible Fitting (MDFF) program operating together with NAMD and VMD (61–63). A scale factor of 1 was used to weigh the contribution of the cryo-EM map to the overall potential function used in MDFF. Simulations included 10,000 steps of minimization and 100,000 steps of molecular dynamics under implicit solvent conditions with secondary structure restraints in place. The refined models were validated using the MolProbity server (64).

Comparison of the maps was done in UCSF Chimera 1.15 by implementing “subtract maps” feature. The maps were calculated at the same resolution and threshold value. Unmodelled densities were identified by creating a density map around the atomic model with the “molmap” command and subtracting it from the corresponding reconstruction.

### Calculations of electrostatic potential of capsid proteins

Final atomic models of capsid proteins of intact and faf-BSA treated CVA9 were used for calculations of electrostatic potential. PDB files were converted to PQR using the PDB2PQR online tool (https://server.poissonboltzmann.org/pdb2pqr) with PROPKA at pH 7.0 to assign protonation states with CHARMM force field and leaving additional options as the default settings. Output results were used for Adaptive Poisson-Boltzmann Solver (APBS) calculations with the Poisson-Boltzmann equation (65). Input parameters were left as the default settings. The output PQR file was used for visualization in ChimeraX 1.3 (66, 67). Surface representation of the input asymmetric subunit PDB file was colored based on the PQR file electrostatic potential values in a range from −10 to 10 from the mean.

### Data availability

The atomic models and cryo-EM maps generated during the current study are available in the wwPDB repositories with accession numbers PDB ID 8AT5, EMD-15634 (CVA9 control), PDB ID 8AW6, EMD-15692 (faf-BSA-treated CVA9) and PDB ID 8AXX, EMD-15706 (ion-treated CVA9).

## Acknowledgements

We thank Pasi Laurinmäki of Instruct-ERIC Centre Finland and the Biocenter Finland National Cryo-Electron Microscopy Unit, Helsinki University, and Saana Haarma, University of Helsinki, for excellent technical assistance. The facilities and expertise of the HiLIFE Biocomplex unit at the University of Helsinki, a member of Instruct-ERIC Centre Finland, FINStruct, and Biocenter Finland are gratefully acknowledged. We thank Antti Tuhkala, Xiaonan Liu and Salla Keskitalo from Proteomics Unit, Institute of Biotechnology and Helsinki Institute of Life Science (HiLIFE), University of Helsinki, Helsinki, Finland, for technical assistance of mass spectrometry analysis. We thank CSC-IT Center for Science Ltd for providing technical assistance and facilities to carry out the work. Molecular graphics and analyses for electrostatic potential were performed with UCSF ChimeraX, developed by the Resource for Biocomputing, Visualization, and Informatics at the University of California, San Francisco, with support from National Institutes of Health R01-GM129325 and the Office of Cyber Infrastructure and Computational Biology, National Institute of Allergy and Infectious Diseases.

This project was supported by the Academy of Finland (grants 315950 and 336471 to SJB), the Sigrid Juselius Foundation (95-7202-38 to SJB), and Jane and Aatos Erkko Foundation (to SJB and VM). ZP is a fellow of the ILS doctoral programme.

## Author contributions

S.J.B. and V.M. conceived the idea. S.J.B., V.M, A.D. designed the experiments. A.D. B.L., Z.P., and V.R. carried out the experiments. A.D., Z.P., B.L. did image analysis. Z.P. and A.D. built the atomic models. V.R. and Z.P prepared the figures. Z.P, V.R., A.D. and S.J.B. interpreted the results. A.D., V.R. and S.J.B. wrote the manuscript. All authors edited and agreed on the final manuscript.

## References

1. Aldriweesh MA, Shafaay EA, Alwatban SM, Alkethami OM, Aljuraisi FN, Bosaeed M, et al. Viruses Causing Aseptic Meningitis: A Tertiary Medical Center Experience With a Multiplex PCR Assay. Front Neurol. 2020;11:602267.

2. Huang YC, Chu YH, Yen TY, Huang WC, Huang LM, Cheng AL, et al. Clinical features and phylogenetic analysis of Coxsackievirus A9 in Northern Taiwan in 2011. BMC Infect Dis. 2013;13:33.

3. Lafolie J, Labbe A, L’Honneur AS, Madhi F, Pereira B, Decobert M, et al. Assessment of blood enterovirus PCR testing in paediatric populations with fever without source, sepsis-like disease, or suspected meningitis: a prospective, multicentre, observational cohort study. Lancet Infect Dis. 2018;18(12):1385–96.

4. Shabani A, Makvandi M, Samarbafzadeh A, Teimoori A, Rasti M, Karami C, et al. Echovirus 30 and coxsackievirus A9 infection among young neonates with sepsis in Iran. Iran J Microbiol. 2018;10(4):258–65.

5. Jiang P, Liu Y, Ma HC, Paul AV, Wimmer E. Picornavirus morphogenesis. Microbiol Mol Biol Rev. 2014;78(3):418–37.

6. Ruokolainen V, Domanska A, Laajala M, Pelliccia M, Butcher SJ, Marjomaki V. Extracellular Albumin and Endosomal Ions Prime Enterovirus Particles for Uncoating That Can Be Prevented by Fatty Acid Saturation. J Virol. 2019;93(17).

7. Zhang P, Mueller S, Morais MC, Bator CM, Bowman VD, Hafenstein S, et al. Crystal structure of CD155 and electron microscopic studies of its complexes with polioviruses. Proc Natl Acad Sci U S A. 2008;105(47):18284–9.

8. Ren J, Wang X, Hu Z, Gao Q, Sun Y, Li X, et al. Picornavirus uncoating intermediate captured in atomic detail. Nat Commun. 2013;4:1929.

9. Liu Y, Sheng J, Baggen J, Meng G, Xiao C, Thibaut HJ, et al. Sialic acid-dependent cell entry of human enterovirus D68. Nat Commun. 2015;6:8865.

10. Strauss M, Schotte L, Karunatilaka KS, Filman DJ, Hogle JM. Cryo-electron Microscopy Structures of Expanded Poliovirus with VHHs Sample the Conformational Repertoire of the Expanded State. J Virol. 2017;91(3).

11. Danthi P, Tosteson M, Li QH, Chow M. Genome delivery and ion channel properties are altered in VP4 mutants of poliovirus. J Virol. 2003;77(9):5266–74.

12. Smyth M, Pettitt T, Symonds A, Martin J. Identification of the pocket factors in a picornavirus. Arch Virol. 2003;148(6):1225–33.

13. Rossmann MG, He Y, Kuhn RJ. Picornavirus-receptor interactions. Trends Microbiol. 2002;10(7):324–31.

14. Milstone AM, Petrella J, Sanchez MD, Mahmud M, Whitbeck JC, Bergelson JM. Interaction with coxsackievirus and adenovirus receptor, but not with decay-accelerating factor (DAF), induces A-particle formation in a DAF-binding coxsackievirus B3 isolate. J Virol. 2005;79(1):655–60.

15. Huttunen M, Waris M, Kajander R, Hyypia T, Marjomaki V. Coxsackievirus A9 infects cells via nonacidic multivesicular bodies. J Virol. 2014;88(9):5138–51.

16. Myllynen M, Kazmertsuk A, Marjomaki V. A Novel Open and Infectious Form of Echovirus 1. J Virol. 2016;90(15):6759–70.

17. Shakeel S, Seitsonen JJ, Kajander T, Laurinmaki P, Hyypia T, Susi P, et al. Structural and functional analysis of coxsackievirus A9 integrin alphavbeta6 binding and uncoating. J Virol. 2013;87(7):3943–51.

18. Karjalainen M, Rintanen N, Lehkonen M, Kallio K, Maki A, Hellstrom K, et al. Echovirus 1 infection depends on biogenesis of novel multivesicular bodies. Cell Microbiol. 2011;13(12):1975–95.

19. Baggen J, Thibaut HJ, Strating J, van Kuppeveld FJM. The life cycle of non-polio enteroviruses and how to target it. Nat Rev Microbiol. 2018;16(6):368–81.

20. Ward T, Powell RM, Chaudhry Y, Meredith J, Almond JW, Kraus W, et al. Fatty acid-depleted albumin induces the formation of echovirus A particles. J Virol. 2000;74(7):3410–2.

21. Carson SD, Cole AJ. Albumin Enhances the Rate at Which Coxsackievirus B3 Strain 28 Converts to A-Particles. J Virol. 2020;94(6).

22. Raiborg C, Bache KG, Gillooly DJ, Madshus IH, Stang E, Stenmark H. Hrs sorts ubiquitinated proteins into clathrin-coated microdomains of early endosomes. Nat Cell Biol. 2002;4(5):394–8.

23. Scott CC, Gruenberg J. Ion flux and the function of endosomes and lysosomes: pH is just the start: the flux of ions across endosomal membranes influences endosome function not only through regulation of the luminal pH. Bioessays. 2011;33(2):103–10.

24. Gerasimenko JV, Tepikin AV, Petersen OH, Gerasimenko OV. Calcium uptake via endocytosis with rapid release from acidifying endosomes. Curr Biol. 1998;8(24):1335–8.

25. Christensen KA, Myers JT, Swanson JA. pH-dependent regulation of lysosomal calcium in macrophages. J Cell Sci. 2002;115(Pt 3):599–607.

26. Steinberg BE, Huynh KK, Brodovitch A, Jabs S, Stauber T, Jentsch TJ, et al. A cation counterflux supports lysosomal acidification. J Cell Biol. 2010;189(7):1171–86.

27. Ruokolainen V, Laajala M, Marjomaki V. Real-time Fluorescence Measurement of Enterovirus Uncoating. Bio Protoc. 2020;10(7):e3582.

28. Curry S, Chow M, Hogle JM. The poliovirus 135S particle is infectious. J Virol. 1996;70(10):7125–31.

29. Hendry E, Hatanaka H, Fry E, Smyth M, Tate J, Stanway G, et al. The crystal structure of coxsackievirus A9: new insights into the uncoating mechanisms of enteroviruses. Structure. 1999;7(12):1527–38.

30. Brandenburg B, Lee LY, Lakadamyali M, Rust MJ, Zhuang X, Hogle JM. Imaging poliovirus entry in live cells. PLoS Biol. 2007;5(7):e183.

31. Brabec M, Schober D, Wagner E, Bayer N, Murphy RF, Blaas D, et al. Opening of size-selective pores in endosomes during human rhinovirus serotype 2 in vivo uncoating monitored by single-organelle flow analysis. J Virol. 2005;79(2):1008–16.

32. Strauss M, Levy HC, Bostina M, Filman DJ, Hogle JM. RNA transfer from poliovirus 135S particles across membranes is mediated by long umbilical connectors. J Virol. 2013;87(7):3903–14.

33. Bilek G, Matscheko NM, Pickl-Herk A, Weiss VU, Subirats X, Kenndler E, et al. Liposomal nanocontainers as models for viral infection: monitoring viral genomic RNA transfer through lipid membranes. J Virol. 2011;85(16):8368–75.

34. Tubiana L, Bozic AL, Micheletti C, Podgornik R. Synonymous mutations reduce genome compactness in icosahedral ssRNA viruses. Biophys J. 2015;108(1):194–202.

35. Petrov AS, Bowman JC, Harvey SC, Williams LD. Bidentate RNA-magnesium clamps: on the origin of the special role of magnesium in RNA folding. RNA. 2011;17(2):291–7.

36. Draper DE. Folding of RNA tertiary structure: Linkages between backbone phosphates, ions, and water. Biopolymers. 2013;99(12):1105–13.

37. Pickl-Herk A, Luque D, Vives-Adrian L, Querol-Audi J, Garriga D, Trus BL, et al. Uncoating of common cold virus is preceded by RNA switching as determined by X-ray and cryo-EM analyses of the subviral A-particle. Proc Natl Acad Sci U S A. 2013;110(50):20063–8.

38. Guo P, Li Y, An J, Shen S, Dou H. Study on structure-function of starch by asymmetrical flow field-flow fractionation coupled with multiple detectors: A review. Carbohydr Polym. 2019;226:115330.

39. Kotoucek J, Hezova R, Vrablikova A, Hubatka F, Kulich P, Macaulay S, et al. Characterization and purification of pentameric chimeric protein particles using asymmetric flow field-flow fractionation coupled with multiple detectors. Anal Bioanal Chem. 2021;413(14):3749–61.

40. Liu X, Salokas K, Weldatsadik RG, Gawriyski L, Varjosalo M. Combined proximity labeling and affinity purification-mass spectrometry workflow for mapping and visualizing protein interaction networks. Nat Protoc. 2020;15(10):3182–211.

41. Meier F, Brunner AD, Koch S, Koch H, Lubeck M, Krause M, et al. Online Parallel Accumulation-Serial Fragmentation (PASEF) with a Novel Trapped Ion Mobility Mass Spectrometer. Mol Cell Proteomics. 2018;17(12):2534–45.

42. Yu F, Haynes SE, Teo GC, Avtonomov DM, Polasky DA, Nesvizhskii AI. Fast Quantitative Analysis of timsTOF PASEF Data with MSFragger and IonQuant. Mol Cell Proteomics. 2020;19(9):1575–85.

43. Chang KH, Auvinen P, Hyypia T, Stanway G. The nucleotide sequence of coxsackievirus A9; implications for receptor binding and enterovirus classification. J Gen Virol. 1989;70 (Pt 12):3269–80.

44. Zheng SQ, Palovcak E, Armache JP, Verba KA, Cheng Y, Agard DA. MotionCor2: anisotropic correction of beam-induced motion for improved cryo-electron microscopy. Nat Methods. 2017;14(4):331–2.

45. de la Rosa-Trevin JM, Quintana A, Del Cano L, Zaldivar A, Foche I, Gutierrez J, et al. Scipion: A software framework toward integration, reproducibility and validation in 3D electron microscopy. J Struct Biol. 2016;195(1):93–9.

46. Rohou A, Grigorieff N. CTFFIND4: Fast and accurate defocus estimation from electron micrographs. J Struct Biol. 2015;192(2):216–21.

47. Mindell JA, Grigorieff N. Accurate determination of local defocus and specimen tilt in electron microscopy. J Struct Biol. 2003;142(3):334–47.

48. de la Rosa-Trevin JM, Oton J, Marabini R, Zaldivar A, Vargas J, Carazo JM, et al. Xmipp 3.0: an improved software suite for image processing in electron microscopy. J Struct Biol. 2013;184(2):321–8.

49. Scheres SH. RELION: implementation of a Bayesian approach to cryo-EM structure determination. J Struct Biol. 2012;180(3):519–30.

50. Abrishami V, Zaldivar-Peraza A, de la Rosa-Trevin JM, Vargas J, Oton J, Marabini R, et al. A pattern matching approach to the automatic selection of particles from low-contrast electron micrographs. Bioinformatics. 2013;29(19):2460–8.

51. Scheres SH. A Bayesian view on cryo-EM structure determination. J Mol Biol. 2012;415(2):406–18.

52. Kimanius D, Forsberg BO, Scheres SH, Lindahl E. Accelerated cryo-EM structure determination with parallelisation using GPUs in RELION-2. Elife. 2016;5.

53. Zivanov J, Nakane T, Forsberg BO, Kimanius D, Hagen WJ, Lindahl E, et al. New tools for automated high-resolution cryo-EM structure determination in RELION-3. Elife. 2018;7.

54. Scheres SH. Beam-induced motion correction for sub-megadalton cryo-EM particles. Elife. 2014;3:e03665.

55. Chen Y, Pfeffer S, Hrabe T, Schuller JM, Forster F. Fast and accurate reference-free alignment of subtomograms. J Struct Biol. 2013;182(3):235–45.

56. Sorzano CO, de la Rosa Trevin JM, Oton J, Vega JJ, Cuenca J, Zaldivar-Peraza A, et al. Semiautomatic, high-throughput, high-resolution protocol for three-dimensional reconstruction of single particles in electron microscopy. Methods Mol Biol. 2013;950:171–93.

57. Rosenthal PB, Henderson R. Optimal determination of particle orientation, absolute hand, and contrast loss in single-particle electron cryomicroscopy. J Mol Biol. 2003;333(4):721–45.

58. Pettersen EF, Goddard TD, Huang CC, Couch GS, Greenblatt DM, Meng EC, et al. UCSF Chimera--a visualization system for exploratory research and analysis. J Comput Chem. 2004;25(13):1605–12.

59. Emsley P, Lohkamp B, Scott WG, Cowtan K. Features and development of Coot. Acta Crystallogr D Biol Crystallogr. 2010;66(Pt 4):486–501.

60. Yang J, Zhang Y. I-TASSER server: new development for protein structure and function predictions. Nucleic Acids Res. 2015;43(W1):W174–81.

61. Phillips JC, Braun R, Wang W, Gumbart J, Tajkhorshid E, Villa E, et al. Scalable molecular dynamics with NAMD. J Comput Chem. 2005;26(16):1781–802.

62. Trabuco LG, Villa E, Mitra K, Frank J, Schulten K. Flexible fitting of atomic structures into electron microscopy maps using molecular dynamics. Structure. 2008;16(5):673–83.

63. Humphrey W, Dalke A, Schulten K. VMD: visual molecular dynamics. J Mol Graph. 1996;14(1):33–8, 27-8.

64. Chen VB, Arendall WB, 3rd, Headd JJ, Keedy DA, Immormino RM, Kapral GJ, et al. MolProbity: all-atom structure validation for macromolecular crystallography. Acta Crystallogr D Biol Crystallogr. 2010;66(Pt 1):12–21.

65. Jurrus E, Engel D, Star K, Monson K, Brandi J, Felberg LE, et al. Improvements to the APBS biomolecular solvation software suite. Protein Sci. 2018;27(1):112–28.

66. Pettersen EF, Goddard TD, Huang CC, Meng EC, Couch GS, Croll TI, et al. UCSF ChimeraX: Structure visualization for researchers, educators, and developers. Protein Sci. 2021;30(1):70–82.

67. Goddard TD, Huang CC, Meng EC, Pettersen EF, Couch GS, Morris JH, et al. UCSF ChimeraX: Meeting modern challenges in visualization and analysis. Protein Sci. 2018;27(1):14–25.

